# StructuralDPPIV: A novel deep learning model based on atom-structure for predicting dipeptidyl peptidase-IV inhibitory peptides

**DOI:** 10.1101/2023.05.22.541389

**Authors:** Ding Wang, Junru Jin, Zhongshen Li, Yu Wang, Mushuang Fan, Sirui Liang, Ran Su, Leyi Wei

## Abstract

**Motivation:** Diabetes is a chronic metabolic disorder that has been a major cause of blindness, kidney failure, heart attacks, stroke, and lower limb amputation across the world. To alleviate the impact of diabetes, researchers have developed the next generation of anti-diabetic drugs, known as dipeptidyl peptidase IV inhibitory peptides (DPP-IV-IPs). However, the discovery of these promising drugs has been restricted due to the lack of effective peptide-mining tools.

**Results:** Here, we presented StructuralDPPI V, a deep learning model designed for DPP-IV-IP identification, which takes advantage of both molecular graph features in amino acid and sequence information. Experimental results on the independent test dataset and two wet experiment datasets show that our model outperforms the other state-of-art methods. Moreover, to better study what StructuralDPPIV learns, we used CAM technology and perturbation experiment to analyze our model, which yielded interpretable insights into the reasoning behind prediction results.

**Availability:** The project code is available at https://github.com/WeiLab-BioChem/Structural-DPP-IV.

## 1 Introduction

Type 2 diabetes is the most common form of diabetes world-wide(Chatterjee, et al., 2017), accounting for 90-95% of all cases of diabetes. It may not be diagnosed until several years after the onset of illness and complications develop. Until recently, this form of diabetes was seen only in adults, but it is now also occurring with increasing frequency in children(Copeland, et al., 2013). Dipeptidyl peptidase IV (DPP-IV, E.C.3.4.14.5) is a cell-surface aminopeptidase that was originally characterized as a T-cell differentiation antigen(Kikkawa, et al., 2006). This enzyme is related to immune regulation, signal transduction and apoptosis(Golightly, et al., 2012). The proteolytic activity of DPP-IV had been classified as a serine-type protease that cleaves a proline or alanine residue at the N-terminal dipeptides position(Casrouge, et al., 2018). DPP-IV inhibitory peptides (DPP-IV-IPs) had been widely considered a promising anti-type 2 diabetes drug(De, et al., 2019). This way of using inhibitory peptides for treatment causes fewer adverse reactions to patients compared with chemically synthesized small-molecule drugs(Barnett, 2006; Jarvis, et al., 2013). Therefore, it is crucial to develop new DPP-IV inhibitory drugs and functional foods.

Traditional laboratory-based techniques for predicting DPP-IV-IP are precise but resource-intensive, expensive, and time-consuming(Nongonierma and FitzGerald, 2019; Wang, et al., 2021). Enzymatic hydrolytic screening is still the main approach used, which hampers the efficiency of DPP-IV-IP screening. With the improvement and supplementation of peptide datasets, several machine learning based predictors have been developed to predict DPP-IV-IP(Charoenkwan, et al., 2020; Guan, et al., 2022; Phasit, et al., 2022). Despite these advancements, these models still exhibit some limitations, and further improvements are possible. iDPPIV-SCM is the first ML-based DPP-IV-IP predictor using scoring card method (SCM), it provided accuracies of 0.819 and 0.797 for cross-validation and independent datasets respectively, but lacks informative features from various facets. This model is not yet accurate enough for real-world applications, and lacked biochemical interpretability(Charoenkwan, et al., 2020). Stack-DPPIV uses combined machine learning methods and utilized more feature encodings from multiple perspectives. It achieved an accuracy of 0.891, but depends on feature engineering for sequence and structure information extracting(Phasit, et al., 2022). BERT-DPPIV(Guan, et al., 2022) is the current state-of-the-art predictor with satisfactory accuracy. However, it only uses NLP method for predicting and depends on time-costing pretraining procedure, which lacks physicochemical information.

In the present study, we proposed a sequence structure-based machine learning predictor (termed StructuralDPP-IV), for DPP-IV-IP identification. This predictor used joint representation extracted from both NLP method (TextCNN) and amino acid structure. To utilize the information from amino acid structure, StructuralDPP-IV used SMILES representation as a medium to extract physicochemical features, which made the model well interpretable from atomic level. Furthermore, on an independent test dataset, as well as the Pentapeptide and Tripeptide wet experiment dataset(Guan, et al., 2022), StructuralDPPIV consistently achieves an accuracy rate surpassing 90%. Notably, it outperforms other established methodologies, thus affirming the robustness and efficacy of our model. To delve deeper into the model’s underlying mechanisms, we employed CAM and permutation analyses to investigate the intricate relationship between predicted peptide sequences, amino acids, and their respective positions. The resulting interpretable findings, of particular interest, align with biological knowledge and statistical scrutiny, thereby offering valuable insights and recommendations for peptide design.

## 2 Materials and Methods

### 2.1 Dataset

We employed Charoenkwan et al.’s dataset(Phasit, et al., 2022) in this study for model development and assessment. The dataset contains 665 positive and 665 negative samples, which is divided into benchmark set for training and test set for independent testing. Benchmark set contains 532 DPP-IV-IPs and 532 negative samples, while independent test set includes 113 DPP-IV-IPs and the same number of non-DPP-IV-IPs.

Pentapeptide and Tripeptide wet experiment dataset is derived from the biological experiment in BERT-DPPIV(Guan, et al., 2022). Pentapeptide dataset consists of peptides that contain proline or alanine and repeat the dipeptide unit VP or IPI. Tripeptide dataset consists of four kind classes, i.e., WPX, WAX, WRX, and WVX, in which X means any possible kind in 20 human amino acids. All the wet biological experiment results and labels are given in the paper of BERT-DPPIV.

### 2.2 StructuralDPP-IV architecture

**Fig.1** shows the StructuralDPPIV workflow and model composition. Our model consists of three parts: (B) Structural encoding module, (C) TextCNN encoding module and (C) Classification module. Given a peptide sequence, we use text convolutional neural module (TextCNN) and structural encoding module to encode representation at sequence level and representation at atom structure level respectively. The process of obtaining a joint representation for a peptide involved two subrepresentations derived from the TextCNN module and the Structural module, respectively. To fuse the information contained in these subrepresentations, we employ an element-wise multiplication of the two-dimensional vectors. The resulting joint representation captures the combined features of the peptide as captured by both the textual and structural information. We subsequently utilize this joint representation to predict whether the peptide is expected to be DPP-IV-IP. This methodology enables the integration of multiple sources of information to enhance the accuracy of predictions regarding the properties of peptides.

**Fig 1.**
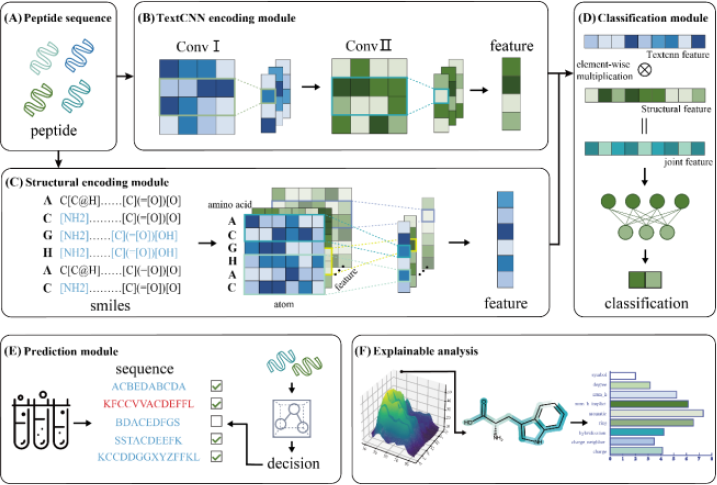
Overview of the StructuralDPP-IV framework. (A) Peptide sequence. We get one sequence from the dataset. (B) TextCNN encoding module. This module is used to encode the peptide from sequence information by using TextCNN model. (C) Structural encoding module. This module is used to encode peptide from the perspective of amino acid structural information. (D) Classification module. After we get two subrepresentations of peptide from TextCNN module and Structural module respectively, we use element-wise multiplication to fuse two-dimensional vector to get the joint representation. Then, we use this joint representation to predict whether this peptide is expected to be DPP-IV-IP. (E) Prediction module. In this work, we verify our model’s validity by utilizing two wet biological experiments. (F) Explainable analysis. We utilize two explainable methods i.e., CAM technology and perturbation experiments to analyze the knowledge learned from our model.

#### 2.2.1 SMILES representation of peptides

Effective representation of molecular structure is a crucial step in the present study. SMILES(simplified molecular input line entry system) is a specification in the form of a line notation for describing the structure of chemical species using short ASCII strings(Weininger, 1988). It had been widely used in many fields of bioinformatics, e.g., interaction and binding affinity prediction of drug–target(Karimi, et al., 2019; Yang, et al., 2021; Zeng, et al., 2021), peptide toxicity prediction(Wei, et al., 2021). Recent studies show that SMILES was used to encode DNA for the prediction of rice 6mA sites by deep learning methods and achieved good performance(Liu, et al., 2022).

#### 2.2.2 Structural encoding module

Structural encoding module encodes peptide’s SMILES representation to get physicochemical information. It is a crucial step to embed the biochemical information contained in SMILES into the features. For one peptide sequence *S* with amino acid *a*_1_, *a*_2_, …, *a*_*n*_ where *n* is the number of atoms in one amino acid molecule, *S*(*a*_1_, *a*_2_, …, *a*_*n*_) is encoded into one 3d tensor ***T***_***S***_ of shape (*n*^*′*^ × *m*^*′*^ × *p*), where *n*^*′*^ is the max sequence length in training and independent test dataset (90 for the dataset from (Charoenkwan, et al., 2020)), *m*^*′*^ is the max number of non-hydrogen atoms in one amino acid (15 for human race) and *p* is the feature number extracted from one atom (21 for our model). Here we note ***T***_***S***_ as ***T***_***S***_(***M***_**1**_, ***M***_**2**_, …, ***M***_***n***_) where ***M***_***n***_ is feature representation matrix of amino acid *a*_*n*_, and ***M***_***i***_ as ***M***_***i***_(***v***_**1**_, ***v***_**2**_, …, ***v***_***m***_) where ***v***_***i***_(*f*_1_, *f*_2_, …, *f*_*p*_) is feature vector of one atom *t*_*i*_. The meanings of these features are shown in the **Table 1**.

**Table 1.** All the 21 features encoded by structural feature extraction part.

| Atom Feature | Contents | Possible Value |
| --- | --- | --- |
| 0, 1, 2, 3 | One-hot-encoding, C, N, O or S atom | True or False |
| 4, 5, 6 | One-hot-encoding, Atom degree | True or False |
| 7, 8, 9, 10 | One-hot-encoding, num of Hs on the atom | True or False |
| 11, 12, 13, 14 | One-hot-encoding, number of implicit Hs on the atom. | True or False |
| 15 | One-hot-encoding, Atom is Aromatic or not | True or False |
| 16, 17 | One-hot-encoding, Atom is in ring or not | 0 or 1 |
| 18 | Atom’s hybridization type | Int, 1-7 |
| 19 | Atom’s Gasteiger charges | Float |
| 20 | Atom’s Gasteiger charges * | Float |

For each atom *t*, feature 0-19 is multiplied with adjacent matrix of this atom. Thus, the value encoded in one entry includes information about neighbor nodes. In the **Table 1**, feature 20 (marked with *) is attached to the multiplication result as one information from one atom itself instead of that from neighbors. Given a peptide sequence, e.g., ‘ACDEFGHIKLMNPQRSTVWY’, Structural encoding module first encodes each char i.e., each amino acid into SMILES sequence representation, then generates one modular object using acquired SMILES with RDKit.

We chose these features based on their biochemical relevance. The atomic features mentioned above comprehensively encapsulate the biochemical characteristics of each individual atom. Since the final vector representation for each atom is a result of the features of neighboring atoms multiplied by the adjacency matrix, this approach effectively conveys physicochemical information for each atom. Through our experimental tests, we found that the inclusion of atomic properties such as atomic number and second-degree neighbor features did not lead to an improvement in model performance.

For encoding features like atomic types, we employed one-hot encoding. This encoding method is advantageous for non-ordinal categorical data as it helps avoid bias and can provide some degree of regularization. Our experiments showed that one-hot encoding was effective in improving model performance when encoding atomic types, whereas for other features, it either slightly outperformed direct encoding or showed minimal differences.

Graph convolutional network is a generalization of convolution on graph structured data and is able to capture complex relationships between nodes in a graph, which has been widely used in many bioinformatic tasks(Chu, et al., 2021; Li, et al., 2020; Ryu, et al., 2020). Compared with direct connection between neural network layers, the residuals block in ResNet is able to effectively prevent gradient from exploding or disappearing in deep network through shortcut connection(He, et al., 2016; He, et al., 2016). In StructuralDPPIV, we utilize residual blocks for further information extraction. The workflow of Structural encoding module is as follows:

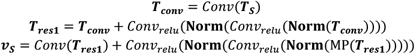

where ***T***_***conv***_, ***T***_***res*1**_ and ***v***_***S***_ are the output of the convolutional layer, the first and the second residual block, respectively. **Norm** denotes batch normalization, MP means max-pooling operation. In Structural encoding module, we first input features of 3d tensor ***T***_***S***_ encoded from peptide sequence into GCN layer, then feed the output into two residual blocks to extract more effective and distinguishable features.

#### 2.2.3 TextCNN encoding module

Text convolutional neural network (TextCNN) utilizes convolutional neural networks (CNN) for sentence-level classification tasks. It utilizes various filter sizes applied to a sentence, capturing diverse latent features. Resulting features are then pooled to generate a fixed-length representation, which is subsequently fed into a linear layer for prediction. TextCNN achieved satisfactory accuracy while maintaining good performance on multiple benchmarks(Kim, 2014). In recent years, TextCNN had been widely used in many bioinformatic fields such as cancer pathology reports classification(Alawad, et al., 2019; Alawad, et al., 2020; Savova, et al., 2019) and bioactive peptide discovery(He, et al., 2022). In our model, we use TextCNN to provide a simple and intuitive NLP representation of peptide sequences. The following equations illustrate how this module works:

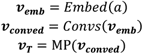

where *a* refers to amino acid, *Embed* refers to embedding layer, *Convs* refers to convolution layer, and MP refers to max-pooling operation. TextCNN encoding module process one amino acid into an embedding vector ***v***_***emb***_, one peptide sequence with *m* acids into tensor ***M***(***v***_**1**_, ***v***_**2**_, …, ***v***_***m***_). In this article, we use representation of sequence with non-static channels i.e., vector embedding of amino acid can be updated during training by backpropagation. Different convolutional filters with weights ***w*** are then applied to ***M***, gaining new embedding ***v***_***conved***_ which is further used for Max-over-time pooling operation. The final output, noted as ***v***_***T***_, of linear layers behind max-pooling module is concatenated with ***v***_***S***_ for classification:

#### 2.2.4 Joint representation

The final stage of the model processing entails element-wise multiplication of the output vectors from the Structural and TextCNN modules. For example, the structural embedding vector (**v**_**S**_) (1, 4, 2, 8) multiplies text embedding vector (**v**_**T**_) (5, 7, 1, 4) get the final representation vector (5, 28, 2, 32). Element-wise multiplication introduces a non-linear interaction between the elements of the two feature vectors. This can capture complex relationships between features that may not be captured by simple addition, and this step is exactly the fusion mechanism mentioned above. The resulting vector is then used in classification task:

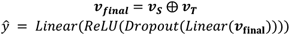

where ***v***_***S***_, ***v***_***T***_ is intermediate representation derived from Structural encoding module or TextCNN encoding module, ***v***_**final**_ is the vector used for classification, ⨁ is element-wise multiplication operation, and Linear, Dropout and ReLU refer to corresponding neural networks.

### 2.3 Evaluation metrics

To evaluate the prediction performance of our model, we utilize five metrics commonly used in binary classification tasks(Charoenkwan, et al., 2021):

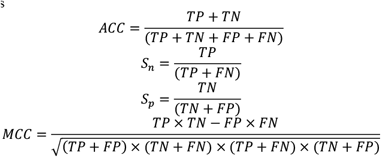

where TP, TN, FP, FN are the number of true positives, true negatives, false positives and false negatives respectively. Sp and Sn stand for Specificity and Sensitivity respectively, they can reflect the model’s ability to recognize DPP-IV inhibitory peptides and non-DPP-IV inhibitory peptides. ACC means Several previous works have employed these metrics to evaluate model performance. Typically, a higher value for ACC, Sn, Sp and MCC metrics indicates better performance of the model.

### Training process

Here we provide comprehensive information about the training process of StructuralDPPIV. Computational calculations were executed utilizing an NVIDIA RTX 3090 GPU equipped with 24 GB of memory. Prior to the training process, preprocessing of each peptide record was performed to derive intermediate structural and textual encodings, herein referred to as intermediate SE and TE tensors. It is noteworthy that SE tensor was directly generated by physicochemical features, and intermediate TE was one-hot dictionary tensor. Subsequently, these tensors were persistently stored on disk as pickle (.pkl) files. This nonparametric preprocessing step yields a significant acceleration in both the subsequent training and testing phases.

The training hyperparameters employed in this study are as follows: a batch size of 32, a learning rate of 0.000005, and a maximum number of epochs set at 150, beyond which the model’s performance reaches a sufficiently stable state. To further enhance performance, we leveraged the Focal Loss(Lin, et al., 2017) and Adam(Kingma and Ba, 2014) optimizer for better performance. The training process consumes approximately 8 minutes.

## 3 Results and discussion

### 3.1 StructuralDPPIV outperforms other State-of-art methods on independent dataset

In this section, we compared StructuralDPPIV with previously reported DPP-IV-IP discriminant models, including Decision Tree(DT) (Dhall, et al., 2021), iDPPIV-SCM(Charoenkwan, et al., 2020), Random Forest(RF) (Charoenkwan, et al., 2021; Dhall, et al., 2021), Naïve Bayes (NB) (Charoenkwan, et al., 2020; Jia and He, 2016), K nearest neighbor(KNN) (Hasan, et al., 2020; Liu and Chen, 2020), fuzzy KNN(fKNN) (Min, et al., 2013), SVM(Zou and Yin, 2021), StackDPPIV(Phasit, et al., 2022) and Bert-DPPIV(Guan, et al., 2022). The experimental results indicated that StructuralDPPIV outperforms other DPP-IV-IP predictors, demonstrating a superior accuracy and MCC of 1.58% and 3.18%, respectively, compared to the current state-of-the-art DPP-IV-IP predictor, BERT-DPPIV. Further comparative results can be found in **Table 2** and **Fig.2 A**. In summary, StructuralDPPIV excels in this comparison due to its deep learning architecture, effectively capturing complex patterns and relationships within peptide sequences. It demonstrates superior classification performance, particularly in terms of high sensitivity and MCC, highlighting its capability to address class imbalance and provide a balanced classification measure. In addition, StructuralDPPIV also demonstrates a high level of interpretability that was not present in previous prediction models. This success stems from its comprehensive and reasonable utilization of physicochemical and sequential information.

**Table 2.** The comparison between StructuralDPPIV and other existing models.

| Model | ACC | MCC | Sn | Sp | AUC |
| --- | --- | --- | --- | --- | --- |
| **StructuralDPP-IV** | 0.9098 | 0.8218 | 0.9474 | 0.8722 | 0.9656 |
| **BERT-DPP-IV** | 0.8940 | 0.7900 | 0.8720 | 0.9170 | 0.9600 |
| **StackDPP-IV** | 0.8910 | 0.7840 | 0.8570 | 0.9250 | 0.9610 |
| **SVM** | 0.8650 | 0.7310 | 0.8740 | 0.8560 | 0.9390 |
| **iDPP-IV-SCM** | 0.7970 | 0.5940 | 0.7890 | 0.8050 | 0.8470 |
| **KNN** | 0.8571 | 0.7163 | 0.8947 | 0.8195 |  |
| **fKNN** | 0.8609 | 0.7243 | 0.9023 | 0.8195 |  |
| **DT** | 0.8722 | 0.7444 | 0.8797 | 0.8647 |  |
| **NB** | 0.8083 | 0.6229 | 0.8797 | 0.7368 |  |
| **RF** | 0.8271 | 0.6589 | 0.8872 | 0.7669 |  |

**Fig 2.**
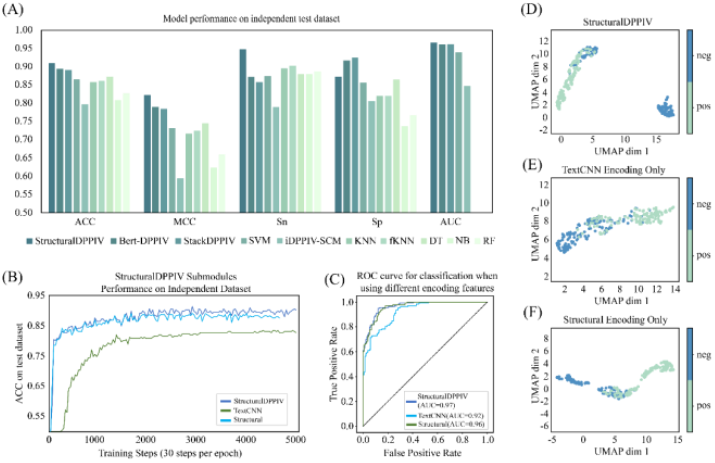
StructuralDPPIV outperforms other models. (A) Different model performances on independent test dataset with different metrics (AUC record of DT, NB, RF is not yet available). (B) The training process of StructuralDPPIV and its submodules. (C) ROC curve of StructuralDPPIV and its submodules. (D-F). Umap analysis of StructuralDPPIV and its submodules. This graph shows StructuralDPPIV has a satisfactory ability to distinguish peptide samples compared with single module.

### 3.2 Ablation experiment identified the importance of submodules in StructuralDPPIV

To demonstrate the predictive mechanism of StructuralDPPIV, we compared the model using only the TextCNN encoding module or only the Structural encoding module with the original model. The experimental results in **Fig.2 B** and **Table 3** indicated that the Structural encoding module played a critical role in the model’s prediction process, with a significant improvement in all metrics. However, the combination of Structural and TextCNN encoding modules enhanced prediction robustness with a more stable training process. On the independent test dataset, the complete model exhibited significantly higher accuracy than that achieved by any single encoding module in isolation.

**Table 3.** The comparison between StructuralDPPIV and other existing models.

| Model | ACC | MCC | Sn | Sp | AUC |
| --- | --- | --- | --- | --- | --- |
| StructuralDPPIV | 0.9098 | 0.8218 | 0.9474 | 0.8722 | 0.9656 |
| TextCNN encoding only | 0.8346 | 0.6869 | 0.9473 | 0.7218 | 0.9226 |
| Structural encoding only | 0.8909 | 0.7918 | 0.9699 | 0.8120 | 0.9629 |

Subsequently, we utilized the Umap(McInnes, et al., 2018) to visualize the output features processed from StructuralDPPIV and submodules. The Umap results demonstrated that StructuralDPPIV has a significantly better ability to distinguish peptide samples than using a single Structural encoding or TextCNN encoding module. As shown in **Fig.2 D**, most negative samples were effectively separated from positive samples in the representation space. In contrast, **Fig.2 E** and **F** showed that the single module cannot project positive and negative examples to different locations in the intermediate representation space, with some negative and positive samples almost connected in the handover area.

### 3.3 The performance of StructuralDPPIV on polypeptide screening

To further verify the reliability of our model, we applied StructuralDPPIV to two other datasets: the Pentapeptide and Tripeptide wet experiment datasets. **Fig.3 A** compared the performance of StructuralDPPIV with BERT-DPPIV. According to the wet experiment conducted by the same author (Guan, et al., 2022), 22 pentapeptides are qualified inhibitory peptides, and BERT-DPPIV predicted 14 of the 22 pentapeptides to exhibit inhibitory activity against DPP-IV. In contrast, StructuralDPPIV successfully predicted 21 of the 22 pentapeptides to be DPP-IV-IPs, with only 4.5% error. On Tripeptide dataset, 72 out of 80 tripeptides were found to efficiently inhibit DPP-IV. We utilized StructuralDPPIV on this dataset and obtained an accuracy of 90%, which is comparable to that of BERT-DPPIV.

**Fig 3.**
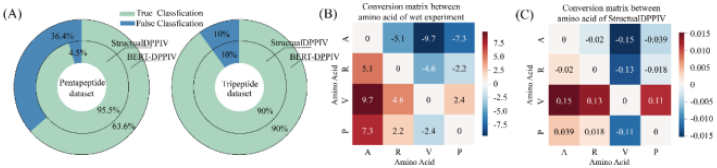
Pentapeptides and Tripeptides screening of StructuralDPPIV, which is previously conducted using BERT-DPPIV model. (A) The comparison between StructuralDPPIV and BERT-DPPIV on Pentapeptide and Tripeptide datasets. (B-C) Conversion matrix between selected amino acids according to wet experiments and StructuralDPPIV.

Furthermore, we investigated the conversion results between different amino acids using wet experiments to test the inhibitory activity changes after one amino acid is mutated into another. To ensure that the samples were sufficient, we selected the four amino acids that appeared most frequently in the Tripeptide dataset. The results are presented in **Fig.3 B**, where we observed a significant increase in inhibitory activity when V(*Valine*) is mutated into A(*Alanine*), while the conversion between V(*Valine*) and P(*Proline*) has little influence on inhibitory activity. We use our model prediction score to calculate the same conversion matrix on Tripeptide dataset, which can verify whether our model could learn this regular pattern. Interestingly, as shown in **Fig.3 C**, the distribution of the conversion matrix calculated by our model was similar to the conversion matrix calculated by wet experiment, demonstrating the excellent learning ability and detection power of our model.

### 3.4 Comprehensive analysis on StructuralDPPIV

In order to further analyze the workflow and interpretability of StructuralDPPIV, we conducted a series of experiments to explore working mechanism of our model, including amino acid position analysis, atomic analysis, amino acid type analysis and feature analysis experiments. Experimental results showed that our model has excellent information-capturing ability.

#### 3.4.1 CAM Analysis

CAM (Class Activation Mapping) analysis is a technique employed to produce visual explanations for decisions made by CNN-based models (Zhou, et al., 2016). This technique enhances the transparency of models by using gradients of any target concept that flow into the final convolutional layer to generate a coarse localization map that highlights significant regions in the image for prediction purposes (Selvaraju, et al., 2017). In our study, we utilized CAM to evaluate the importance of different amino acids and atoms in the decision-making process of our model. The CAM score map of amino acid position and atom index is presented in **Fig.4 A**.

**Fig 4.**
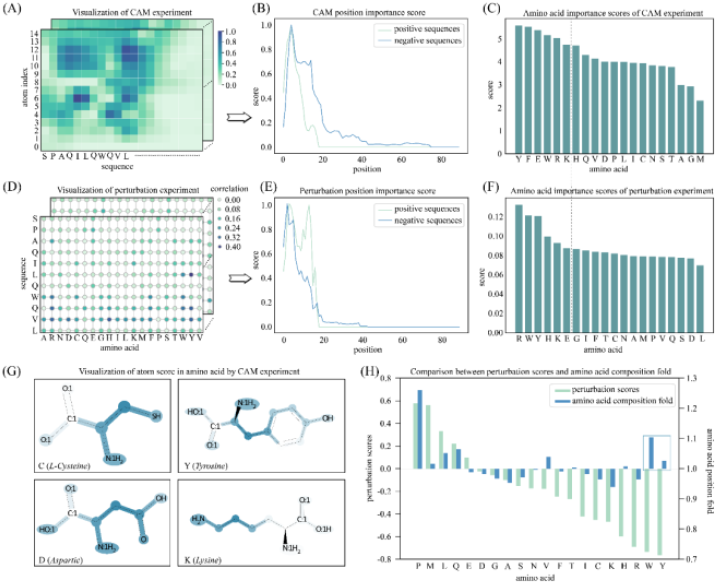
Position and atom importance analysis. (A-C) Class Activation Mapping (CAM) and inspection of amino acid position and amino acid type. (D-F) Perturbation inspection of position and amino acid type. (G) Examples of CAM analysis score map on three amino acids. Experimental results showed that StructuralDPPIV focuses on different R groups of amino acid, therefore making credible decisions. (H) Effect of different amino acid types on prediction.

To make a more comprehensive analysis, we calculated the CAM score of each amino acid statistically for each position of each sequence in the dataset, focusing on position and amino acid type. As demonstrated in **Fig.4 B**, both positive and negative examples have relatively high importance scores before the 14th position, indicating the importance of the head region when designing DPP-IV-IP. We also analyzed the importance score on different amino acid types by adding up CAM score of each amino acid respectively, as illustrated in **Fig.4 C**. Our results indicated that Y(*Tyrosine*), F(*Phenylalanine*), and E(*Glutamic acid*) had the highest importance score, which was useful for DPP-IV-IP designing.

Moreover, CAM analysis was applied to amino acid molecular structure analysis, and the importance score on atom index is presented in **Fig.4 G** (the remained analysis of other amino acids can be seen in the supplementary file). Our analysis of 20 amino acid demonstrated that for most amino acid molecules, StructuralDPPIV mainly focuses on the structure of R groups, instead of paying much attention to amino and carboxyl groups. This was reasonable and consistent with expectations.

#### 3.4.2 Perturbation analysis

Perturbation analysis is a process that involves the replacement of an amino acid in a polypeptide sequence with another amino acid, followed by the recording of the model’s prediction change for the new sequence. Through this analysis, statistical calculation of perturbation score for each amino acid could be executed for each position of each sequence.

It is noteworthy that perturbation analysis also provided results on amino acid position and type that were similar to those obtained using CAM analysis. Interestingly, the results on position importance and amino acid importance were comparable between perturbation experiment and CAM analysis. Specifically, as observed in the CAM analysis on position importance (**Fig.4 E**), the perturbation experiment gave greater attention to the head region, particularly the 4th and 13th positions, which was highly consistent with the CAM analysis. Furthermore, the amino acid type importance score calculated by the perturbation experiment was presented in **Fig. 4 F**. As a gray line runs through the two charts, the left part highly overlapped between perturbation experiment and CAM analysis, sharing the same amino acids Y(*Tyrosine)*, E(*Glutamic acid*), W(*Tryptophan*), R(*Arginine*), and K(*Lysine*). Although the order between them was not entirely consistent, there was an 83% coincidence rate in the composition of the left part.

In the above perturbation analysis, we employed the absolute value to represent the importance score. However, it is still unknown whether a mutation has a positive or negative influence on the prediction score. To address this, we utilized subtractive values. Positive values represent a positive effect, while negative values indicate a negative impact. Furthermore, for statistical verification, we introduce K-fold on AAC (Amino Acid Composition), using one amino acid composition value on positive sequences to divide that on negative sequences. In the realm of AAC fold metrics, it could be observed that the median value is 1. Furthermore, it is worth noting that AAC metrics that exceed 1 demonstrate amino acids exhibiting positive effects, while those yielding negative effects exhibit AAC metrics below 1. The experimental results are shown in **Fig.4 H**. Most amino acid effects are consistent between perturbation scores and AAC fold, which proved the data mining ability of our model. What’s more, in the blue frame, we found that Y(*Tyrosine)* and W(*Tryptophan*) were not consistent between two analyses. So, StructuralDPPIV may have the potential to learn the latent effect of amino acid which can not be notified by general statistics methods.

#### 3.4.3 Feature Encoding Analysis

In this study, we aimed to investigate the relative contribution of different features to the performance of our StructuralDPPIV model, both theoretically and experimentally. So we conducted a set of experiments in which we systematically removed one of the nine feature encodings and recorded model performances under the same conditions. Specifically, we compared the average accuracy rate of models without different feature encodings after training for more than 100 epochs. As shown in **Fig.5 A**, when one feature encoding was removed, the performance of our model decreased to varying degrees, demonstrating the importance of each feature in the model’s prediction process.

**Fig 5.**
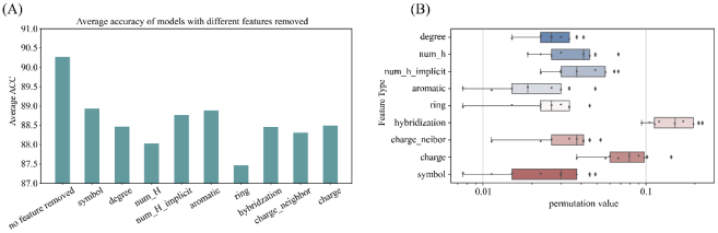
Feature encoding importance analysis. (A) Model performance in circumstances with specific feature encoding removed, compared with the original StructuralDPPIV model. (B) Permutation analysis of different features, which is roughly consistent with the experimental verification results.

To further examine which feature encoding played a more critical role in the prediction accuracy of our model, we utilized permutation importance analysis. This technique measures the decrease in a model score when a single feature value is randomly shuffled (Breiman, 2001). As displayed in **Fig.5 B**, we observed that the hybridization feature had the greatest impact on our model’s performance. However, we acknowledged that the reason for this could be that hybridization is represented by successive values, which may cause the model to be more confused than other feature encodings using discrete values. Additionally, we observed that charge also represents a vital feature encoding, which significantly affects predictive outcomes. It can be postulated that charge information carries implications regarding the polarity of the modules, thereby rendering it a critical property for accurate predictions.

#### 3.4.4 Prospects for Advancement

The dataset involved in this study contains peptides with varying lengths from 2 to 90. Considering the significant variance in sequence lengths, our model employs a padding mechanism to standardize the shape of tensors across different records. We hypothesize that uniform sequence lengths will yield improved performance for StructuralDPPIV. However, in parallel, we have experimented with various padding methods for data augmentation, aiming to mitigate the impact of diverse sequence lengths on model prediction accuracy, but there was no significant improvement observed. Nevertheless, we still believe that better padding or data augmentation methods can further enhance the model’s prediction accuracy. A similar model architecture may exhibit enhanced performance on other datasets containing peptides, DNA, and RNA sequences of uniform lengths. This presents a potential avenue for future research endeavors.

## 4 Conclusions

Recent studies reported that DPP-IV inhibitory peptides (DPP-IV-IP) play a vital role in diabetes study, it is widely used as novel antidiabetic agents to improve glycemic regulation in Type 2 diabetics (T2D) by inhibiting incretin hormones degradation. In the current research, we present StructuralDPPIV as a novel sequence structure-based DPP-IV-IP predictor, which utilizes TextCNN and SMILES representation of peptide sequence and achieved state-of-the-art accuracy on independent test set. The model consists of two modules: Structural encoding module and TextCNN encoding module. Structural Encoding module encodes 21 selected physicochemical features extracted from amino acid, which can be gained from SMILES representation of peptide sequence, then GCN and ResNet are used to extract hidden information and transform input tensor into vector, which will be element-wise multiplied with encoding vector extracted from TextCNN and then used for classification. This model outperformed both TextCNN and Structural single module, and achieved satisfactory results on additional tripeptide and pentapeptide data set. Finally, we conducted explainable analysis experiments, including atomic, feature, position and amino acid type importance analysis. The result shows that StructuralDPPIV is a promising and novel model for DPP-IV-IP prediction.

## Availability of Data and Materials

All code used in data analysis and preparation of the manuscript, alongside a description of necessary steps for reproducing results, can be found in a GitHub repository accompanying this manuscript: https://github.com/WeiLab-BioChem/Structural-DPP-IV.

